# The Cten signalling pathway stabilises Src protein to promote Epithelial-Mesenchymal Transition (EMT) in colorectal cancer

**DOI:** 10.1101/458109

**Authors:** Abdulaziz Asiri, Michael S Toss, Teresa Pereira Raposo, Maham Akhlaq, Hannah Thorpe, Abdulaziz Alfahed, Abutaleb Asiri, Mohammad Ilyas

## Abstract

Cten is an oncogene which promotes epithelial-mesenchymal transition (EMT) in many signalling pathways. Having previously shown that Cten promotes EMT through Snail, we investigated whether Cten function could be mediated through Src (a known regulator of Snail).

Cten levels were modulated by forced expression in colorectal cancer (CRC) cell lines with low Cten expression (HCT116 and RKO) and gene knockdown in a cell line with high Cten expression (SW620). In all cell lines, Cten was a positive regulator of Src expression. The functional importance of Src was tested by forcibly expressing Cten and simultaneously knocking down Src. This resulted in abrogation of Cten motility-inducing activity (cell migration, cell invasion, wound healing – each p<0.001) and abrogation of the promotion of colony formation by Cten (p<0.001) together with failure to induce the Cten targets - Snail and ROCK1. To complement these experiments, Cten expression was restored by forced expression in a subclone of SW620 in which the Cten gene had been deleted (SW620^ΔCten^). SW620^ΔCten^ showed reduced expression of Src which increased following restoration of Cten by forced expression. In SW620^ΔCten^, restoration of Cten increased cell motility (cell migration, cell invasion, wound healing) and colony formation (each p<0.001) which were all lost if Src was concomitantly knocked down. Quantitative Reverse-Transcription PCR (qRT-PCR) showed that modulation of Cten had no effect on Src mRNA levels. However, a cycloheximide (CHX) pulse chase assay demonstrated stabilisation of Src protein by Cten. Finally, the expression of Cten and Src was tested in a series of 84 primary CRCs and there was significant correlation between Cten and Src expression (p=0.001).

We conclude that Src is a novel and functionally important target of the Cten signalling pathway and that Cten protein causes post-transcriptional stabilisation of Src protein in order to promote EMT and possibly metastasis in CRC.

## Introduction

C-terminal tensin-like (Cten, also known as tensin 4) is a member of the tensin protein family which localises to focal adhesion and comprises Tensin 1, Tensin 2, Tensin 3 and Cten/Tensin 4. Tensin 1-3 have extensive sequence and structural homology and have in common an actin binding domain, a Src homology 2 (SH2) domain and a phosphotyrosine binding (PTB) domain. Cten contains SH2 and PTB domains but it lacks the actin binding domain. Tensins localise to focal adhesions and absence of the capability to bind actin filaments is thought to be essential to the role of Cten [1. The importance of Cten is underscored by data showing that it is a common and functionally active target for a multitude of different signalling pathways [2–6].

During tumour invasion, epithelial cells lose their cell to cell adhesion and undergo a shift to a mesenchymal phenotype [7] known as Epithelial-Mesenchymal Transition (EMT) which is thought to be important in promoting cell migration and invasion [8]. EMT has also been characterised by the acquisition of features of stemness [9] and is therefore considered to be a process which contributes to the development of metastasis.

We have previously shown that Cten is involved in the acquisition of fibroblastic features, enhancement of motility (migration and invasion) and enhancement of colony formation efficiency in soft agar and can therefore be regarded as an inducer of EMT [10]. This is mediated in part through proteins such as Integrin-Linked Kinase (ILK) and Focal Adhesion Kinase (FAK) [11,12] and culminates in the up-regulation of Snail and down-regulation of E-cadherin [3,10]. Additionally, we have shown that Cten expression has been associated with metastasis [11]. Although some of the mechanisms of Cten activity have been previously described, the full understanding of additional layers of complexity and signalling pathways that may cooperate with Cten to regulate EMT remain to be investigated.

Src is a non-receptor cytoplasmic tyrosine kinase protein and one of the important kinases in focal adhesion complexes [13]. Src has been found to form a complex with FAK at focal adhesions [12,14]. Src signalling has also been reported to be implicated in regulation of EMT and it has been shown that suppression of Src activity with inhibitor or siRNA knockdown leads to reversal of EMT with up-regulation of E-cadherin and down-regulation of vimentin in breast carcinoma cell lines [15]. Although Src is a well described oncogene, it is rarely mutated in epithelial tumours. This would imply that any role it has in epithelial neoplasia is most probably as a mediator of other oncogenic signalling pathways.

We and others have shown that Cten confers the features which constitute EMT [3,10]. Here we show, for the first time, that Cten signals through Src pathway to increase cell motility and promote colony formation. Furthermore, we demonstrate that Cten is positively correlated with both ROCK1 and Src protein expression in primary colorectal cancers.

## Materials and methods

### Cell Culture

This work was performed in CRC cell lines RKO, HCT116 and SW620, which were a kind gift from Prof Ian Tomlinson. We deleted Cten in SW620 to create the SW620^ΔCten^ cell line as previously described [3]. All cells were cultured in Dulbecco’ s Modified Eagle’ s Medium (DMEM) (Thermo Fisher Scientific, Carlsbad, CA) antibiotic free supplemented with 2mM L-glutamine and 10% foetal bovine serum (FBS) (Sigma, St.Louis, MO) and maintained at 37°C in a 5% CO2 atmosphere. Cell line identities were authenticated by STR profiling.

### Cell Transfection

Lipofectamine 2000 (Thermo Fisher Scientific) was used to transfect the target cells with plasmid DNA and small interfering RNA (siRNA) duplexes following the manufacturer’s protocol. For gene knockdown experiments, cells were grown to 40-50% confluency and 10nM siRNA duplexes targeting Cten, Src or Luciferase (supplementary Table 1) were transfected with 10 μl Lipofectamine. For forced expression experiments, cells were grown to 60-70% confluency and 5 μg the GFP-Cten expression construct [6], or an empty vector control expressing GFP only (GFP-EV), were transfected with 10 μl Lipofectamine in Opti-MEM media (Thermo Fisher Scientific) according to the manufacturer’ s protocol. For co-transfection experiments, cells were grown to 50% confluency and 10 μl Lipofectamine was transfected together with 5 μg plasmid and 10 nM siRNA in Opti-MEM media according to manufacturer’ s instructions. Cells were incubated with the transfection reagents for 6 hours and experimentation performed 48 hours post transfection.

### Cycloheximide Chase Assay

The CHX assay was commenced twenty-four hours post cell transfection. The media in all wells was replaced with fresh DMEM (supplemented with 10% FBS) containing 100 μg/ml CHX. Protein lysates were collected from the cells at different time points ranging from 0–24 hours after CHX exposure. Western blot and densitometry were used to quantify protein and each sample was normalised to β-actin.

### Western Blot

Cell lysates were prepared using RIPA lysis buffer (Thermo Fisher Scientific) supplemented with phosphatase and protease inhibitor (Thermo Fisher Scientific). Fifty microgram of protein was added to NUPAGE LDS Sample Buffer (Thermo Fisher Scientific) containing 5% β-mercaptoethanol. The protein samples were heated to 95 °C on a heat block for 5 minutes and cooled on ice for another 5 minutes. Following this, protein samples were fractionated on a pre-cast 4–12% NUPAGE Bis-Tris-HCl buffered (pH 6.4) polyacrylamide gel (Thermo Fisher Scientific) using the NUPAGE gel electrophoresis system with NUPAGE MOPS SDS Running Buffer (Thermo Fisher Scientific) at 125 V for 90 minutes. Proteins were transferred onto PVDF membrane (GE Life Sciences) using the Trans Blot semi-dry transfer system (Biorad). Following blocking in 5% milk or 5% BSA in 0.1% tween PBS or 0.1% tween TBS (dependent on antibody diluents), membranes were incubated with optimally diluted primary antibodies overnight at 4°C. Following washing, membranes were incubated with the appropriate anti-mouse or anti-rabbit secondary antibody for 1 hour at room temperature (supplementary Table 2). The ECL prime detection kit (GE Life Sciences) was used for protein band visualisation using the C-DiGiT Blot Scanner (LI-COR, Lincoln, NE). Densitometric analysis of the bands was performed using ImageJ gel analysis plugin. Pixel counts for each protein of interest were normalized to β-actin.

### Co-immunoprecipitation

Co-immunoprecipitation (co-IP) was performed to investigate complex formation and binding interaction between proteins. To prepare the cell lysates, a pre-clearing stage was performed to reduce non-specific binding. First, 1 mg of protein was pre-cleared by incubation with 20 μl of Protein G/A agarose beads (Calbiochem IP05) with gentle rotation at 4°C for 30 minutes. The lysate was then centrifuged at 4°C at 13,000 rpm for 5 minutes to pellet the beads and the supernatant retained. Subsequently, 1 μg of ROCK1 (10 μl), Snail (4 μl), Cten (2 μl), or Src (3 μl) antibody was added to 300 μL of the pre-cleared lysate and incubated with gentle rotation overnight at 4°C. In addition, 300 μg of pre-cleared lysate (without antibody) was also incubated with the precleared beads for the negative control. The following day, 30 μl of the Protein A/G beads were added to the reactions and incubated on a rotator for a further 24 hours at 4°C. The beads were pelleted by centrifugation at 13,000 rpm at 4°C for 5 minutes and washed twice in ice cold PBS.

The sample was re-suspended in 30 μl 2X NUPAGE loading Buffer and heated at 95° for 5 minutes, kept on ice for 5 minutes and centrifuged for 2 minutes at 13,000 rpm before loading onto an SDS gel for western blot analysis.

### Quantitative reverse transcription-PCR (qRT-PCR)

qRT-PCR was used to quantify mRNA expression. RNA was extracted using the Total RNA Extraction Kit (Sigma) according to manufacturer’ s protocol. For cDNA synthesis, 1 μg RNA plus 0.5 μg Random hexamers (Thermo Fisher Scientific) were heated for 5 min at 70°C and then cooled for 5 min at 4°C. This was reverse transcribed using 200U of M-MLV Reverse Transcriptase (Promega) and 0.5 mM dNTPs (Promega), heated at 37°C for 1 h followed by 95°C for 10 minutes. Gene quantification was performed on the 7500 Fast Real-Time PCR System (Applied Biosystems) using Go Taq Mastermix (Promega, Madison, WI) and 250 nM each primer (Supplementary Table 3). The run cycle was comprised of a denaturation step at 95°C for 2 minutes, 40 cycles (95°C for 30 seconds, annealing at 60°C for 30 seconds) and a melt curve stage 95°C for 15 seconds, 60°C for 1 minute, 95°C for 15 seconds, and 60°C for 15 seconds. Negative controls included no template and no-RT (RNA only) templates. Each sample was run in triplicate. HPRT was used as the endogenous control and the primer efficiency was determined using a standard curve method. Since primer pairs had similar efficiency the 2^-ΔΔCt^ method was used for gene quantification.

### Transwell Migration and Invasion Assays

The changes in cell migration were assessed in 24 well plates using the Transwell system (Corning, Corning, NY). The Transwell inserts (6.5 mm diameter; 8 μm pore size) were incubated in DMEM at 37°C for 1 hour prior to use. Following this, 600 μl of DMEM (20% FBS) was added to the receiver wells of the Transwell plate and the Transwell inserts placed inside. A total of 1 × 10^5^ cells in DMEM (10% FBS) were seeded onto the Transwell insert. The plate was incubated at 37°C for 24 hours. Following this, the cells that had migrated through and were present on the bottom of the well were manually counted. The Transwell invasion assay was performed similarly with the exception that 2 × 10^5^ cells were seeded onto a Transwell insert coated with growth factor reduced (GFR) Matrigel (0.3 mg/ml, Corning) and cells allowed to migrate for 48 hours prior to counting. Triplicate wells were seeded for each experimental condition.

### Wound Healing

The wound healing ‘scratch assay’ was performed to confirm and assess cell migration. Briefly, 5 to 7 × 10^4^ cells were seeded onto both sides of a culture insert (ibidi) attached in 6 well plates and incubated until confluence (24 hours). After cell attachment, the “scratch” was created by removing the culture inserts, and pictures were taken at various timepoints using an inverted microscope (Nikon) at 10x magnification. The width of the cell free gap was approximately 500 microns (+/− 50 microns) at 0 hours. The wound area was quantified using the MRI Wound Healing Tool (http://dev.mri.cnrs.fr/projects/imagej-macros/wiki/Wound_Healing_Tool), with the ImageJ software. Experiments were performed in duplicate and on at least two separate occasions

### Immunohistochemistry (IHC)

Immunohistochemical (IHC) staining was performed on a tissue microarray slides (TMA) from a series of 84 primary colorectal cancers. This series was created specifically for the purpose of evaluating biomarker expression in colorectal tumours and there was an unbiased selection of consecutive cases collected in Nottingham Sciences Biobank at the Queens Medical Centre, Nottingham, UK. Local ethical approval was granted prior to the construction of the TMAs, destined to IHC studies. Formalin-fixed paraffin embedded tissue blocks were retrieved from the archives and a TMA established as previously described [16].

Immunohistochemical staining was carried out on tissue microarray (TMA) sections (4 μm thickness) mounted on glass slides using the Novolink Polymer Detection Kit (Leica) in accordance with the manufacturer’ s recommended instructions. Briefly, TMA slides were heated at 60°C for10 minutes, followed by dewaxing in xylene and dehydration in industrial methylated spirits (IMS) using the Autostainer XL (Leica). Antigen retrieval was accomplished by heating the samples at 95°C in 10 mM sodium citrate buffer at pH 6.0 or in Tris/EDTA buffer at pH 9.0 for 20 minutes. Slides were then incubated with Peroxidase Block solution at room temperature for 5 minutes and washed with Tris-Buffered Saline (TBS) (pH 7.6) followed by Protein Block for 5 minutes and a TBS wash. Following blocking, the primary antibodies diluted in Antibody Diluent (Leica) were incubated with slides for 1 hour at room temperature (Supplementary Table 4). Slides were then incubated with Post Primary Block for 30 minutes at room temperature. Following this, slides were washed, and Novolink Polymer was applied for 30 minutes, followed by a further wash with TBS. Visualisation of bound antibody was achieved using diaminobenzidine (DAB) prepared from DAB Chromogen and DAB Substrate Buffer. This was incubated for 5 minutes, washed with TBS and following this, slides were counterstained with haematoxylin for 6 minutes. Finally, slides were cleared in xylene and rehydrated in IMS using the Autostainer XL before mounting in DPX.

Whole slides were scanned using the Nanozoomer digital slide scanner (Hamamatsu) and viewed with NDP.view 2 software (Hamamatsu). Immunostaining was scored by AA and MT. Protein expression within the tumour tissues was assessed using the semi-quantitative Histo-score (H-score) where different staining intensities were multiplied by the percentage of representative cells in the tissue for each intensity (0 for negative staining, 1 for weak staining, 2 for moderate staining and 3 for strong staining), producing a range of values between 0 and 300 [17]. Folded cores, those containing exclusively normal and/or stromal tissue or showing tumour tissue less than 15% of the whole core area were excluded from the scoring.

### Statistical Analysis

All *in vitro* experiments were performed using GraphPad Prism (version 6). Results were tested for a normal distribution, and unpaired t-test or the analysis of variance (ANOVA) statistical tests were applied following for experiments with two or more treatment groups respectively.

All statistical analysis of IHC TMA staining was performed using IBM SPSS statistics software (v 22). The expression of protein was categorised into low and high based on the median. The Chi square test was used to test for associations between marker expression (i.e. low or high) and clinicopathological parameters. The Spearman’s rank test was performed to determine correlations between expression of two different markers. For all statistical tests, a p-value < 0.05 was considered significant.

## Results

### Cten is a positive regulator of Src expression in CRC cells

CRC cell lines, HCT116 and RKO which express very low endogenous levels of Cten were transfected with either GFP-EV control (expressing GFP) or GFP-Cten (expressing GFP-tagged Cten) expression constructs. Changes in protein level of Src, ROCK1 and Snail were determined by western blot and quantified by densitometry (Figure 1). We have previously shown that Snail is a target of Cten [3] and, more recently, we have shown that ROCK1 is a target of Cten (Asiri et al., submitted). These constructs were used to validate the effect of modulation of Cten expression.

**Figure 1.**
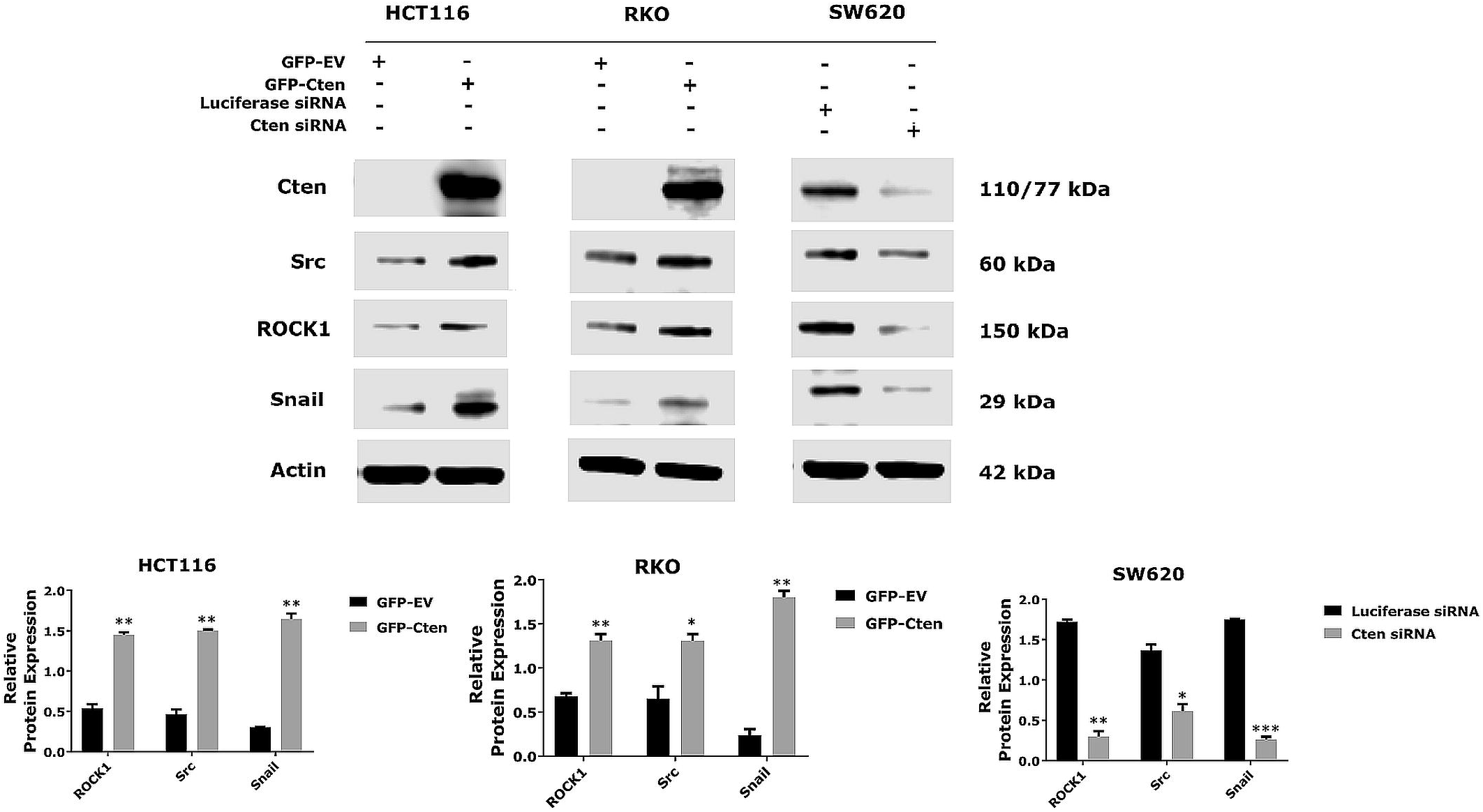
Cten regulates ROCK1, Src, and Snail protein expression. Forced expression of Cten in HCT116 and RKO was associated with an increase in ROCK1, Src, and Snail expression. Knockdown of Cten resulted in a decrease in ROCK1, Src, and Snail protein expression. Graphs on the lower panel represent the densitometry values calculated for each protein band normalised to actin. Results are representative of at least 2 experimental replicates (* = p <0.05, ** = p <0.01, *** = p <0.001).

Forced expression of GFP-Cten in both HCT116 and RKO resulted in upregulation of Src expression as well as ROCK1 and Snail expression compared to cells transfected with GFP-EV control. To confirm this, Cten was also transiently knocked down using Cten targeting siRNA duplexes in SW620 which expresses high endogenous levels of Cten. Knockdown of Cten resulted in downregulation of Src, ROCK1 and Snail protein expression (Figure 1).

### Cten signals through Src to promote motility and colony formation

We next tried to ascertain whether the effects of Cten signalling on Src were functionally relevant to Cten signalling. Cten was forcibly expressed in HCT116 and Src was simultaneously knocked down. When GFP-Cten and luciferase siRNA were co-transfected into HCT116, as expected, there was an increase in Src, ROCK1 and Snail protein levels compared to empty vector (GFP-EV) cotransfected with luciferase siRNA control (Figure 2A). However, when GFP-Cten and Src siRNA were co-transfected (producing a condition of high Cten but depleted Src), the increase in Snail and ROCK1 expression was reduced (Figure 2 A) demonstrating that Src was important in the induction of these Cten targets. The functional importance of Src in Cten signalling was confirmed in a variety of assays. Depletion of Src when Cten was forcibly expressed resulted in a reduction of cell migration (by transwell migration and would healing, p<0.001 for each, Figure 2 B, C) and cell invasion through matrigel (p<0.001, Figure 2 D). We have previously shown that Cten enhances colony formation in soft agar [3] and we observed that Src depletion resulted in a reduction in colony forming efficiency (p<0.001, Figure 2 E).

**Figure 2.**
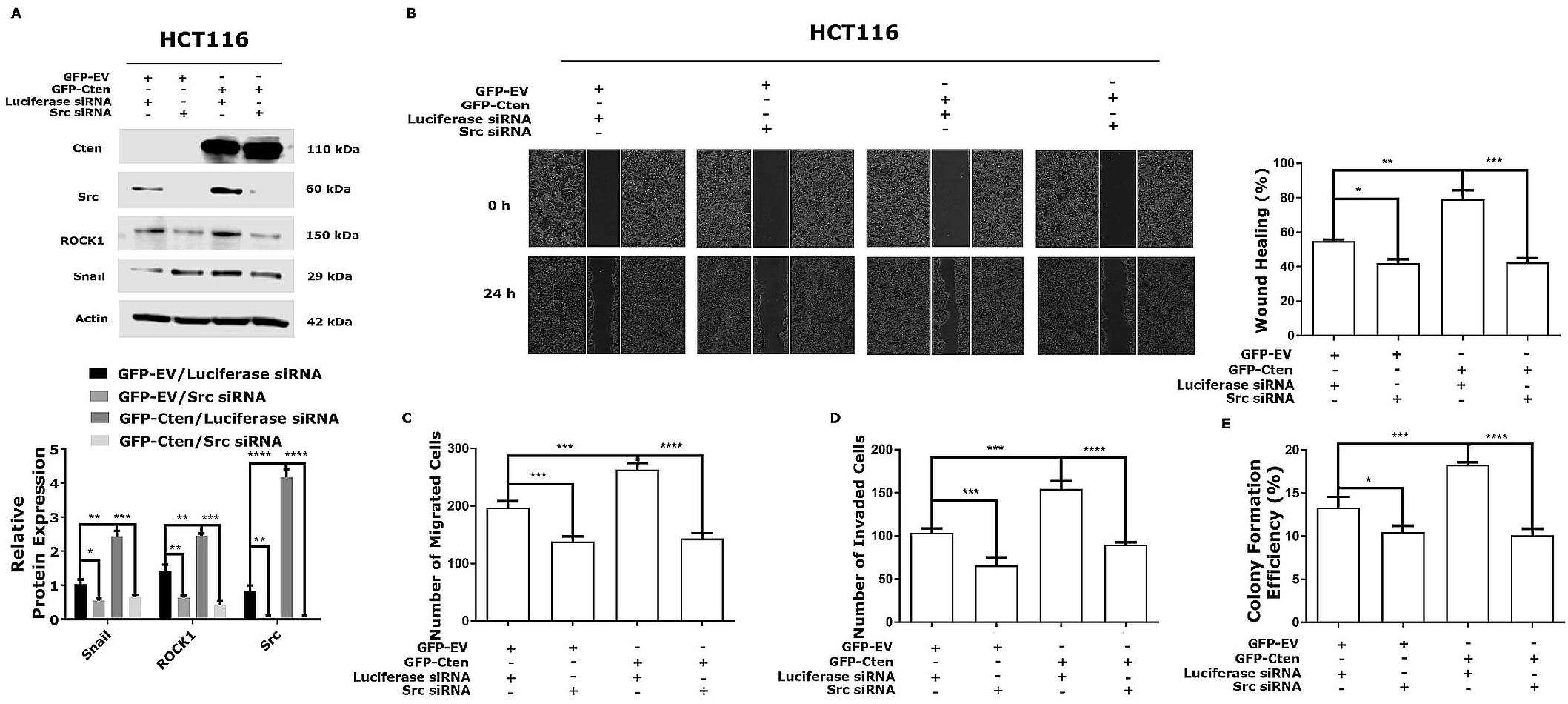
Cten regulates cell functions through Src signalling in HCT116 cells. A) HCT116 cells were overexpressed with GFP-EV or GFP-Cten together with either luciferase or Src targeting siRNA and the changes in Src, ROCK1 and Snail protein expression were determined by western blot. Graph on the lower panel represents the densitometry values calculated for each protein band normalised to actin. B) Wound healing assay showed increased closure of wound following Cten forced expression (P = 0.0049) and this was inhibited when Src was subsequently knocked down (P = 0.0010). C) Overexpression of Cten was associated with an increase in cell migration (P = 0.0002) and this was reduced following Src knockdown (P ≤ 0.0001). D) Overexpression of Cten increased cell invasion (P = 0.0001) and this was lost with subsequent Src knockdown (P ≤ 0.0001). E) Overexpression of Cten induced colony formation efficiency (P = 0.0004) and this was lost on Src knockdown (P ≤ 0.0001). Results are representative of at least 3 experimental replicates (* = p <0.05, ** = p <0.01, *** = p <0.001, **** = p <0.0001).

### Validation of the relationship between Cten and Src

New technologies such as CRISPR allow genes to be somatically deleted and therefore enable protein function to be tested using a variation of Henle-Koch postulates [18]. We have previously described the cell line SW620^ΔCten^ – a subclone and isogenic variant of SW620 in which the Cten gene has been deleted [3]. SW620^ΔCten^ showed lower expression of Src, ROCK1 and Snail protein levels compared to native SW620 (Figure 3 A). Cten was then forcibly re-introduced by transfection of GFP-Cten vector which resulted in the restitution of Src, ROCK1 and Snail protein expression (Figure 3 B). Taken together, the data validate our observations that Cten positively regulates Src and confirm ROCK1 and Snail as targets of Cten.

**Figure 3.**
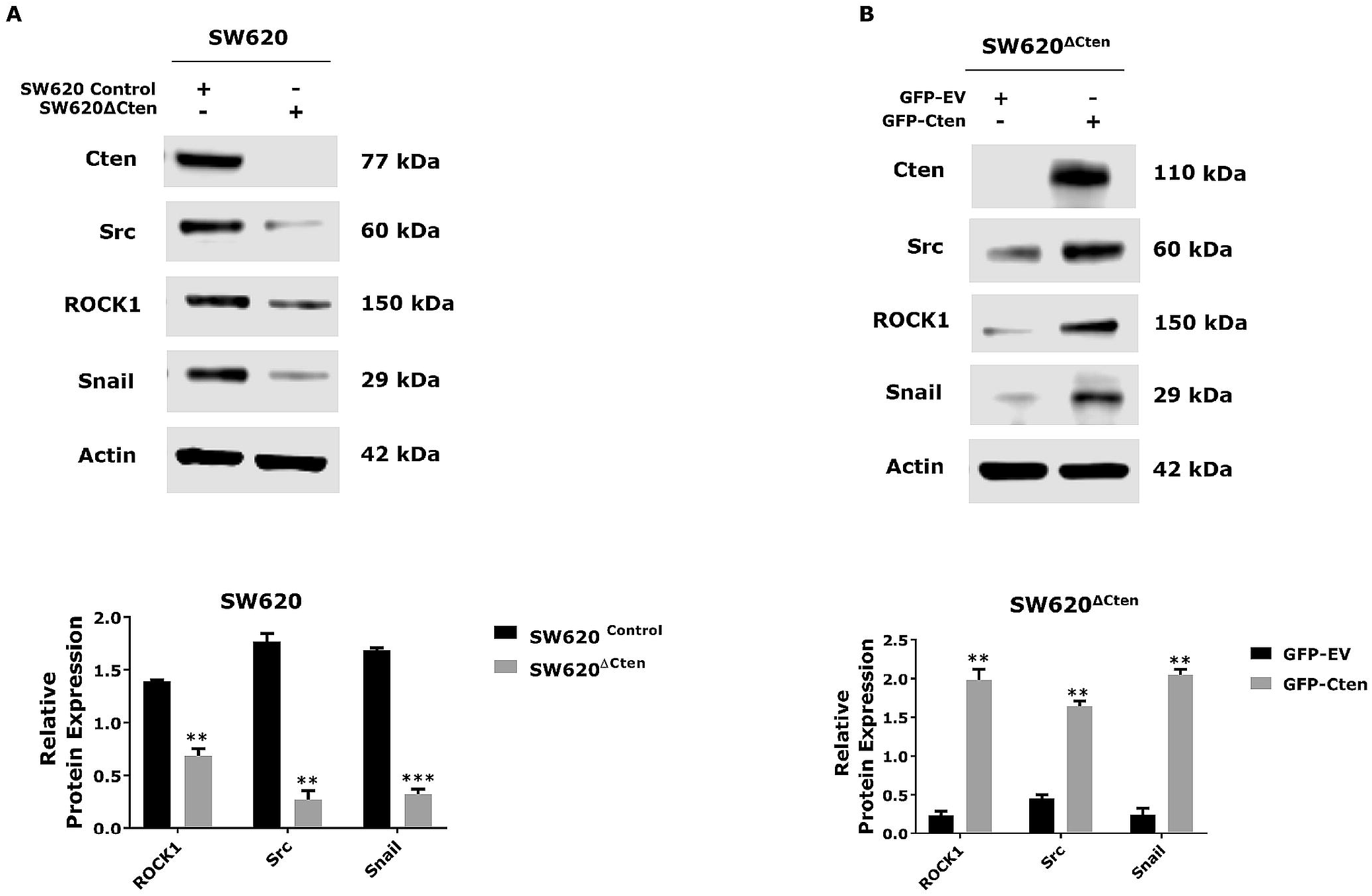
Cten regulates ROCK1, Src, and Snail protein expression. A) Knockout of Cten resulted in a decrease in ROCK1, Src, and Snail protein expression. B) Forced expression of Cten in SW620^ΔCten^ was associated with an increase in ROCK1, Src, and Snail expression. Graphs on the lower panel represent the densitometry values calculated for each protein band normalised to actin. Results are representative of at least 2 experimental replicates (* = p <0.05, ** = p <0.01, *** = p <0.001, **** = p <0.0001).

Functional experiments were repeated in SW620^ΔCten^ i.e. Cten was forcibly re-expressed in SW620^ΔCten^ with concomitant knockdown of Src (Figure 4A). Consistently with previous observations, restitution of Cten expression in SW620^ΔCten^ resulted in up-regulation of Src, Snail and ROCK1 but concomitant depletion of Src resulted in failure to up-regulate Snail and ROCK1. Similarly, restitution of Cten expression in SW620^ΔCten^ resulted in up-regulation of cell motility assessed by wound healing - p<0.05, transwell migration – p<0.01 and invasion – p<0.05 (Figure 4 B, C,D) and colony forming efficiency (p<0.001) (Figure 4 E) when compared to SW620^ΔCten^ transfected with control vector. Depletion of Src and Cten forced expression, resulted in a comparative failure to induce these functions (wound healing (p=0.003), transwell migration, invasion and colony formation (each p<0.001, Figure 4 B, C. D, E).

**Figure 4.**
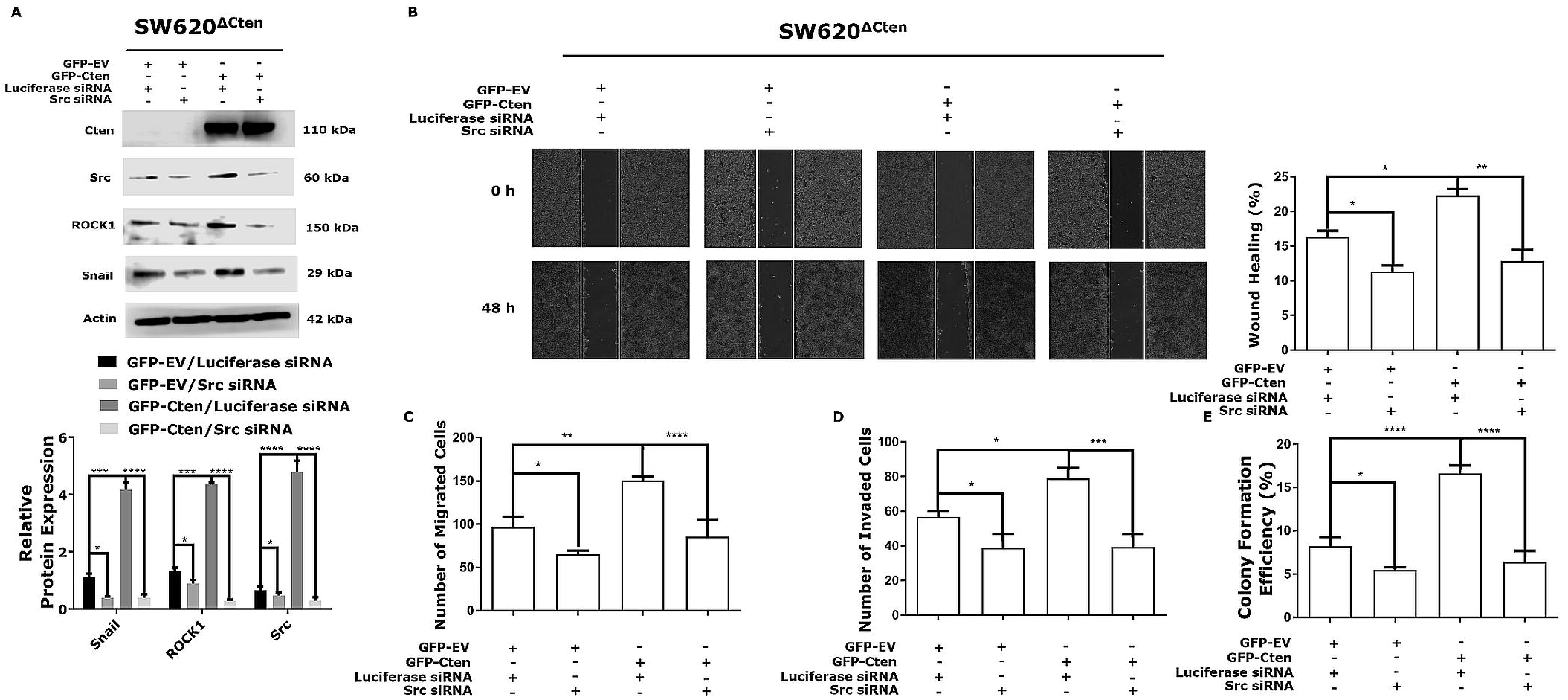
Cten regulates cell functions through Src signalling in SW620^ΔCten^ cells. A) SW620^ΔCten^ cells were overexpressed with GFP-EV or GFP-Cten together with either luciferase or Src targeting siRNA and the changes in Src, ROCK1 and Snail protein expression were determined by western blot. Graph on the lower panel represents the densitometry values calculated for each protein band normalised to actin. B) Overexpression of Cten in SW620^ΔCten^ increased closure of wound (P = 0.0182) which was lost with subsequent Src knockdown (P = 0.0033). C) Overexpression of Cten resulted in an induction of cell migration (P = 0.0021) and this was inhibited when Src subsequently knocked down (P = 0.0006). D) Overexpression of Cten increased cell invasion (P = 0.0124) and this was lost with subsequent Src knockdown (P = 0.0003). E) Overexpression of Cten was associated with an increase in colony formation efficiency (P ≤ 0.0001).and this was lost on Src knockdown (P ≤0.0001). Results are representative of at least 3 experimental replicates (* = p <0.05, ** = p <0.01, *** = p <0.001, **** = p <0.0001).

Taken together, these data suggest Src is an important mediator of Cten signalling upstream of ROCK1 and Snail.

### Cten up-regulates Src expression through protein stabilisation

Having shown that Cten positively regulates Src expression we investigated whether this regulation was at the transcriptional level. Cten was forcibly expressed in HCT116 and knocked down in SW620 cells and quantitative reverse-transcription PCR (qRT-PCR) was performed to quantify changes in Src mRNA expression level. In both cases there was no change in Src mRNA level compared to the control (Figure 5 A, B) indicating that Cten does not induce Src transcription. To investigate whether Cten up-regulates Src expression through post-transcriptional mechanisms such a protein stabilisation, HCT116 cells were transfected with GFP-Cten (or GFP-EV) and treated with CHX. Src protein levels were measured and quantified by densitometry at several time points by Western blot (Figure 5 C and D). The Src decay curve was found to be much steeper (indicating faster decay) in the samples from GFP-EV control than the samples from GFP-Cten transfected cells. The half-life of Src protein was ∼6 h in the cells transfected with GFP-EV control but was appreciably longer, ∼12 h, in the presence of GFP-Cten. This indicates that Cten up-regulates Src protein expression through enhanced protein stability.

**Figure 5.**
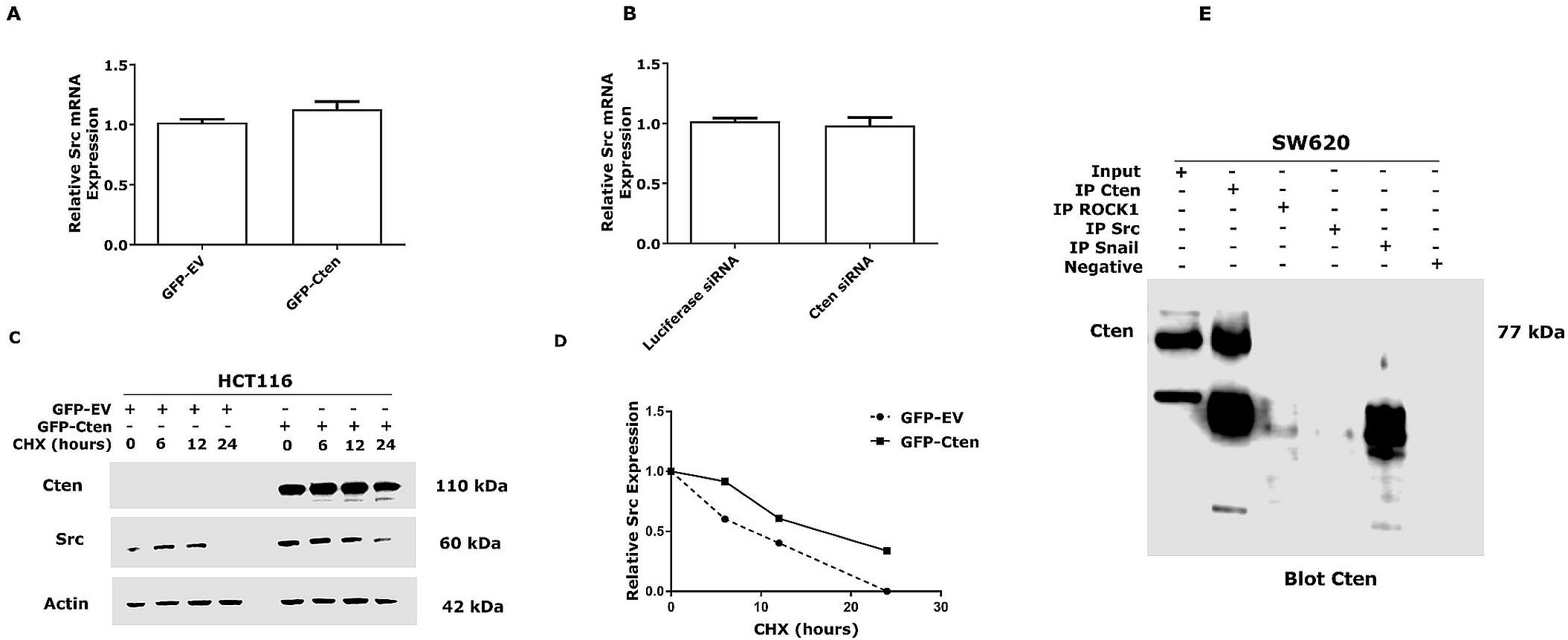
Cten increases Src protein stability. A) Src mRNA expression level did not change following transfection of GFP-Cten in HCT116 cells compared to empty vector (EV) control transfected cells (P = 0.0589). B) The mRNA level of Src expression remined unchanged following transfection of Cten targeting siRNA in SW620 cells compared to luciferase siRNA control transfected cells (P = 0.4696). C) Src protein was stabilised for much longer in HCT116 cells expressing GFP-Cten construct compared to GFP-EV control following treatment with CHX (100 μg/ml) for 0-24 h. D) The Src decay curve following treatment with CHX. Src protein expression was normalised to actin and then normalised to the 0 h time point. E) Endogenous full length Cten does not co-immunoprecipitate with either Src, ROCK1 or Snail. in SW620 cells. Results are representative of at least 3 experimental replicates

Since Cten has been shown to form a complex with a proteins such as β catenin [19], we hypothesised that Cten could form a complex with Src as a potential mechanism for stabilisation of Src protein. Immunoprecipitation (IP) was performed using an antibody to Src but it did not pull down Cten. Similarly, there was no evidence of complex formation with ROCK1, Src or Snail (Figure 5 E).

### Cten, Src, and ROCK1 Expression in Colorectal Cancer

The expression of Cten, Src and ROCK1 was tested in a series of primary CRC cases which was studied using immunohistochemical staining.

Cten expression was shown to localise mostly to the cytoplasm as previously described [10] (Figure 6 A). Cten expression was not associated with the clinical features including tumour grade, tumour stage, lymph node stage, vascular invasion, Dukes’ stage resection margin and KRAS mutation (supplementary Table 5). Both cytoplasmic and membranous expression of Src was seen in the tumour cells (Figure 6B) and each was scored separately. Src staining was not associated with the clinicopathological features as determined using the Chi-square statistical test (supplementary Table 6).

**Figure 6.**
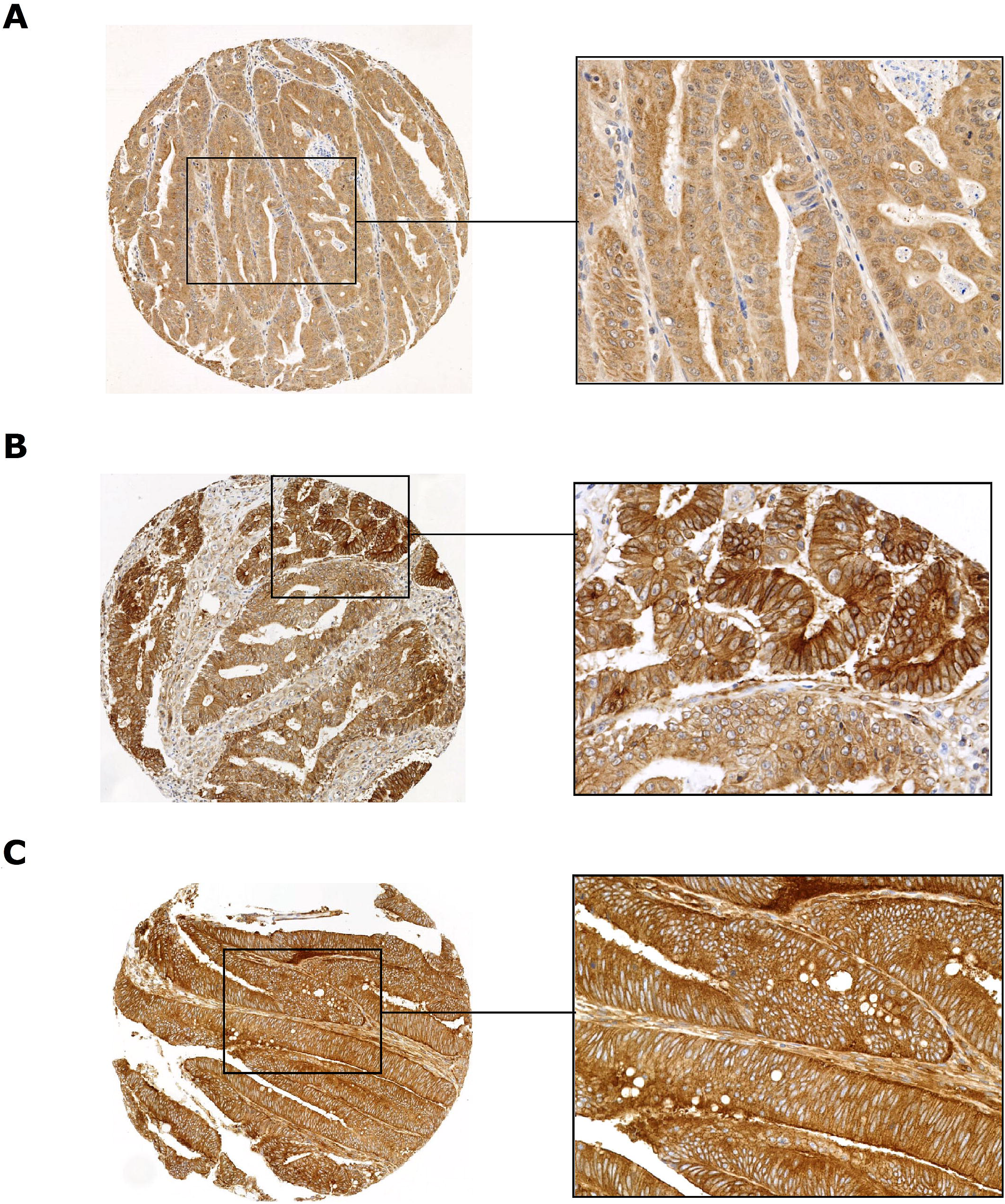
Cten Src, and ROCK1 staining of colorectal tumour tissues. A) TMA cores of CRC revealed Cten staining in the cytoplasm. B) Src staining in the cytoplasm and membrane. C) ROCK1 expression in the cytoplasm.

Comparison of Cten expression and Src expression showed a positive correlation of Cten expression and membranous Src expression (p=0.001) as determined by the Spearman’s rank statistical test (Figure 7 A).

**Figure 7.**
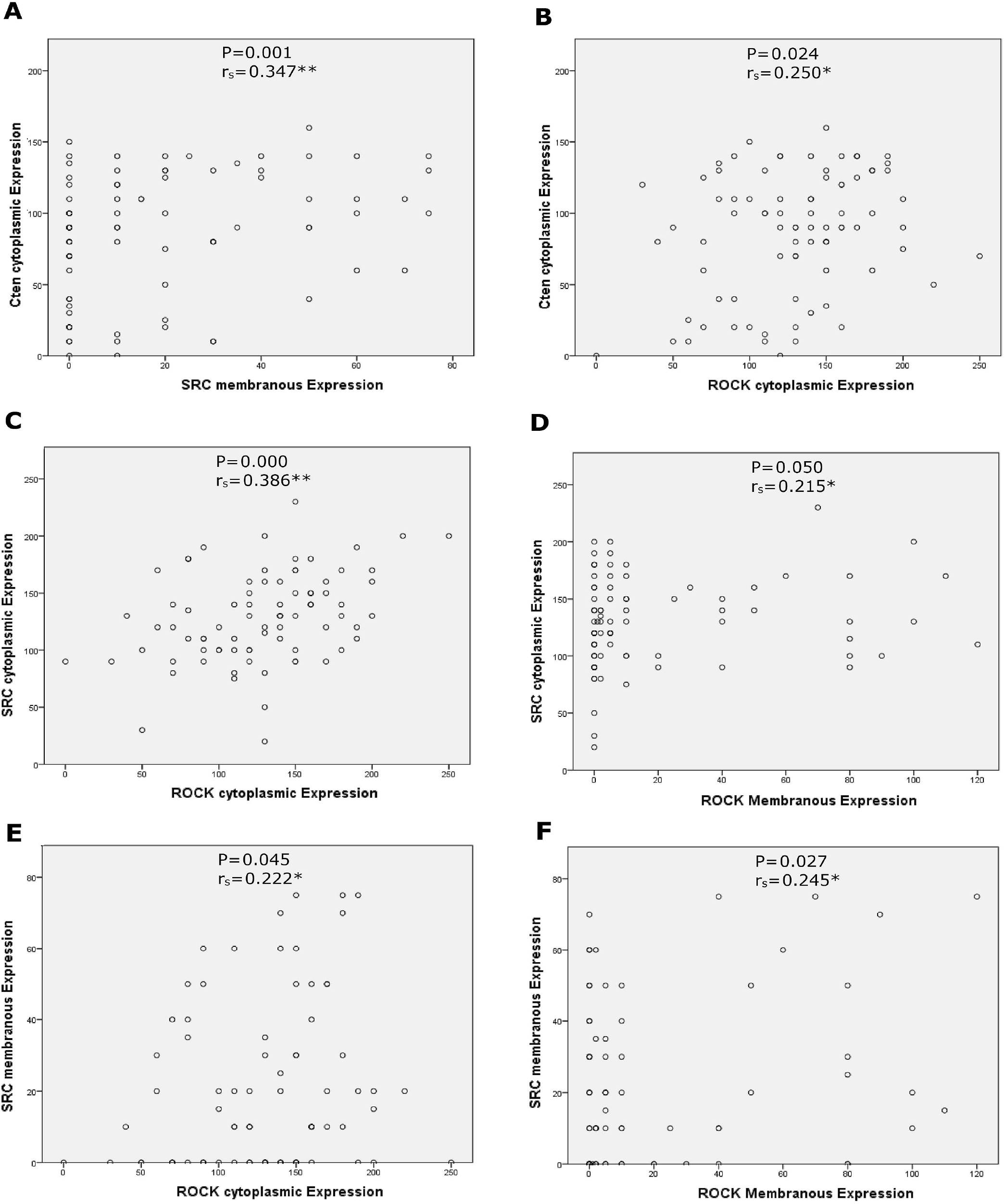
The correlation of Cten and its downstream targets staining of colorectal tumours (Spearman’s rank test): A) Cten cytoplasmic and Src membranous expression (P = 0.001), B) Cten cytoplasmic and ROCK1 cytoplasmic expression (P = 0.024); The correlation of Src and ROCK1 staining of colorectal tumours (Spearman’s rank test). C) The graphs show the correlations of Src cytoplasmic and ROCK1 cytoplasmic (P < 0.001), D) Src cytoplasmic and ROCK1 membranous (P = 0.050), E) Src membranous and ROCK1 cytoplasmic (P = 0.045), F) Src membranous and ROCK1 membranous (P = 0.027).

We have found that ROCK1 is a target of Cten (Asiri et al., submitted) and it would seem that Src is able to induce ROCK1 expression. Immunostaining for ROCK1 was performed and cytoplasmic ROCK1 expression was observed (Figure 6C). ROCK1 staining was not associated with any clinicopathological features (supplementary Table 7). ROCK1 cytoplasmic did however positively correlate with Cten cytoplasmic staining (Figure 7 B) and both ROCK1 cytoplasmic and membranous expression correlate with Src cytoplasmic and membranous staining (Figure 7C,D,E,F).

## Discussion

Cten has been shown to induce EMT in a variety of different tumour tissues [10,12,20–22]. Based on our previous data on the interaction between Cten and Snail [3] and on published data on the interaction between Src and Snail [13,14], we sought to investigate the possibility of Src acting as a target of the Cten signalling pathway. This is the first report of Src as a functionally important target of Cten signalling. Using multiple approaches we have shown that Src is positively regulated by Cten and that, in the absence of Src, the ability of Cten to induce target genes (ROCK1 and Snail) and EMT-related functions (cell motility and colony formation) is reduced. We tried to fulfil the Henle-Koch postulates of causation by demonstrating that deleting Cten in SW620^ΔCten^ results in reduction of EMT features and target gene expression. Ectopic expression of Cten restores these functions/targets and if Src is knocked down, the ability of ectopic Cten to restore the functions/targets is lost.

Although this is the first description of a relationship between Cten and Src, interaction between the two molecules is not surprising since both localise to focal adhesions and Src is reported to be important in the biology of focal adhesion complexes [13,14]. We have provided some mechanistic insight into Cten-Src signalling by showing that upregulation of Src by Cten is due to stabilisation of Src protein rather than transcriptional induction. We investigated whether the mechanisms of stabilisation could be formation of a complex which could stabilise Src. However, immunoprecipitation studies failed to show any physical complex formation between Cten and Src. The mechanism of Cten-mediated Src protein stabilisation is therefore uncertain although it may involve complex formation with other proteins at the focal adhesions or, if it becomes phosphorylated as part of Cten signalling, this may result in stabilisation of the protein [23,24]. Additionally, previous studies have suggested that Src protein stabilisation is most likely to be induced through the upregulation of Akt/mTOR/4E-BP1 pathways and inhibition of calpain-mediated Src protein degradation by ErbB2 signalling[25]. Cten has been shown to be induced by ErbB2 signalling [4] and negatively regulated by calpain [6] One can conjecture that the ErbB2 pathway could protect Cten through calpain inhibition and this in turn could lead to Src protein stabilisation. Therefore, it would be interesting to explore whether Cten signalling is involved in this process.

We tested the expression of Cten, Src and ROCK1 in a series of primary CRCs by immunohistochemistry (IHC). We found that there was a significant positive correlation between Cten expression and Src membranous expression and between Cten and ROCK1 cytoplasmic expression. Furthermore, there was a significant positive correlation between Src and ROCK1. Whilst IHC is a snapshot view of a tumour, these data are supportive of our *in-vitro* observations and our hypothesis that there may be a Cten-Src signalling pathway which then activates ROCK1 and Snail in CRCs. We have previously shown that Cten expression is associated with metastasis [11]. The metastatic process requires acquisition of features of cell migration, cell invasion and “stemness”. Our proposed Cten signalling pathway promotes all of these features and thus supports the possibility that Cten is important in the development of metastasis.

## Conclusions

In summary, we have previously shown that Cten induces EMT and herein we show that this may be mediated through the recruitment of Src. This pathway results in the stabilisation of Src although its mechanism is not clear. This pathway also activates ROCK1 and Snail which may be downstream effectors of the pathway. Further investigations are required to obtain a greater granularity level of this pathway and determine whether these findings could be further explored for therapeutic development of cancer metastasis.

## Acknowledgments

This work was supported by the Saudi Arabian Cultural Bureau in Britain (S3555).

## Author Contributions

AA conducted all the experimental work, analysed the data and wrote the manuscript. MT verified the scoring of the TMAs and assisted with the analysis of the data. TPR reviewed and edited the manuscript. AA and AA assisted with the analysis of the data. MA assisted with the design of the primer sequences and edited the manuscript. HT participated in the TMA staining. MI designed the study and reviewed the manuscript. All authors read and approved the final manuscript.

## References

[1] S.H. Lo, T. Bin Lo, Cten, a COOH-terminal tensin-like protein with prostate restricted expression, is down-regulated in prostate cancer., Cancer Res. 62 (2002) 4217–21.

[2] S. Al-Ghamdi, A. Albasri, J. Cachat, S. Ibrahem, B.A. Muhammad, D. Jackson, A.S. Nateri, K.B. Kindle, M. Ilyas, Cten Is Targeted by Kras Signalling to Regulate Cell Motility in the Colon and Pancreas, PLoS One. 6 (2011) e20919. doi:10.1371/journal.pone.0020919.

[3] H. Thorpe, A. Asiri, M. Akhlaq, M. Ilyas, Cten promotes epithelial-mesenchymal transition through the post-transcriptional stabilization of Snail, Mol. Carcinog. 56 (2017) 2601–2609. doi:10.1002/mc.22704.

[4] M. Katz, I. Amit, A. Citri, T. Shay, S. Carvalho, S. Lavi, F. Milanezi, L. Lyass, N. Amariglio, J. Jacob-Hirsch, N. Ben-Chetrit, G. Tarcic, M. Lindzen, R. Avraham, Y.C. Liao, P. Trusk, A. Lyass, G. Rechavi, N.L. Spector, S.H. Lo, F. Schmitt, S.S. Bacus, Y. Yarden, A reciprocal tensin-3-cten switch mediates EGF-driven mammary cell migration, Nat. Cell Biol. (2007). doi:10.1038/ncb1622.

[5] S.-Y. Hung, Y.-P. Shih, M. Chen, S.H. Lo, Up-regulated cten by FGF2 contributes to FGF2-mediated cell migration, Mol. Carcinog. 53 (2014) 787–792. doi:10.1002/mc.22034.

[6] H. Thorpe, M. Akhlaq, D. Jackson, S. Al Ghamdi, S. Storr, S. Martin, M. Ilyas, Multiple pathways regulate Cten in colorectal cancer without a Tensin switch, Int. J. Exp. Pathol. (2015). doi:10.1111/iep.12154.

[7] S.A. Mani, W. Guo, M.-J. Liao, E.N. Eaton, A. Ayyanan, A.Y. Zhou, M. Brooks, F. Reinhard, C.C. Zhang, M. Shipitsin, L.L. Campbell, K. Polyak, C. Brisken, J. Yang, R.A. Weinberg, The Epithelial-Mesenchymal Transition Generates Cells with Properties of Stem Cells, Cell. 133 (2008) 704–715. doi:10.1016/j.cell.2008.03.027.

[8] K.T. Yeung, J. Yang, Epithelial-mesenchymal transition in tumor metastasis, Mol. Oncol. 11 (2017) 28–39. doi:10.1002/1878-0261.12017.

[9] C.D. May, N. Sphyris, K.W. Evans, S.J. Werden, W. Guo, S.A. Mani, Epithelialmesenchymal transition and cancer stem cells: a dangerously dynamic duo in breast cancer progression., Breast Cancer Res. 13 (2011) 202. doi:10.1186/bcr2789.

[10] A. Albasri, R. Seth, D. Jackson, A. Benhasouna, S. Crook, A.S. Nateri, R. Chapman, M. Ilyas, C-terminal Tensin-like (CTEN) is an oncogene which alters cell motility possibly through repression of E-cadherin in colorectal cancer, J. Pathol. (2009). doi:10.1002/path.2508.

[11] A. Albasri, S. Al-Ghamdi, W. Fadhil, M. Aleskandarany, Y.C. Liao, D. Jackson, D.N. Lobo, S.H. Lo, R. Kumari, L. Durrant, S. Watson, K.B. Kindle, M. Ilyas, Cten signals through integrin-linked kinase (ILK) and may promote metastasis in colorectal cancer, Oncogene. (2011). doi:10.1038/onc.2011.26.

[12] S. Al-Ghamdi, J. Cachat, A. Albasri, M. Ahmed, D. Jackson, A. Zaitoun, N. Guppy, W.R. Otto, M.R. Alison, K.B. Kindle, M. Ilyas, C-Terminal Tensin-Like Gene Functions as an Oncogene and Promotes Cell Motility in Pancreatic Cancer, Pancreas. 42 (2013) 135–140. doi:10.1097/MPA.0b013e3182557ceb.

[13] E. Ingley, Src family kinases: Regulation of their activities, levels and identification of new pathways, Biochim. Biophys. Acta - Proteins Proteomics. 1784 (2008) 56–65. doi:10.1016/j.bbapap.2007.08.012.

[14] V. Bolós, J.M. Gasent, S. López-Tarruella, E. Grande, The dual kinase complex FAK-Src as a promising therapeutic target in cancer., Onco. Targets. Ther. 3 (2010) 83–97.

[15] X. Liu, R. Feng, Inhibition of epithelial to mesenchymal transition in metastatic breast carcinoma cells by c-Src suppression., Acta Biochim. Biophys. Sin. (Shanghai). 42 (2010) 496–501. doi:10.1093/abbs/gmq043.

[16] A. Albasri, W. Fadhil, J.H. Scholefield, L.G. Durrant, M. Ilyas, Nuclear expression of phosphorylated focal adhesion kinase is associated with poor prognosis in human colorectal cancer., Anticancer Res. 34 (2014) 3969–74.

[17] M. Shehata, A. Mukherjee, S. Deen, A. Al-Attar, L.G. Durrant, S. Chan, Human leukocyte antigen class I expression is an independent prognostic factor in advanced ovarian cancer resistant to first-line platinum chemotherapy, Br. J. Cancer. 101 (2009) 1321–1328. doi:10.1038/sj.bjc.6605315.

[18] F. Zhang, Y. Wen, X. Guo, CRISPR/Cas9 for genome editing: progress, implications and challenges, Hum. Mol. Genet. 23 (2014) R40–R46. doi:10.1093/hmg/ddu125.

[19] Y.C. Liao, N.T. Chen, Y.P. Shih, Y. Dong, H. Lo Su, Up-regulation of C-terminal tensin-like molecule promotes the tumorigenicity of colon cancer through β-catenin, Cancer Res. (2009). doi:10.1158/0008-5472.CAN-09-0117.

[20] H. Sasaki, S. Moriyama, K. Mizuno, H. Yukiue, A. Konishi, M. Yano, M. Kaji, I. Fukai, M. Kiriyama, Y. Yamakawa, Y. Fujii, Cten mRNA expression was correlated with tumor progression in lung cancers., Lung Cancer. 40 (2003) 151–5.

[21] H. Sasaki, H. Yukiue, Y. Kobayashi, I. Fukai, Y. Fujii, Cten mRNA Expression Is Correlated with Tumor Progression in Thymoma, Tumor Biol. 24 (2003) 271–274. doi:10.1159/000076141.

[22] A. Albasri, M. Aleskandarany, A. Benhasouna, D.G. Powe, I.O. Ellis, M. Ilyas, A.R. Green, CTEN (C-terminal tensin-like), a novel oncogene overexpressed in invasive breast carcinoma of poor prognosis, Breast Cancer Res. Treat. (2011). doi:10.1007/s10549-010-0890-3.

[23] R. Sears, F. Nuckolls, E. Haura, Y. Taya, K. Tamai, J.R. Nevins, Multiple Ras-dependent phosphorylation pathways regulate Myc protein stability., Genes Dev. 14 (2000) 2501–14.

[24] V. Lopez-Pajares, M.M. Kim, Z.-M. Yuan, Phosphorylation of MDMX Mediated by Akt Leads to Stabilization and Induces 14-3-3 Binding, J. Biol. Chem. 283 (2008) 13707–13713. doi:10.1074/jbc.M710030200.

[25] M. Tan, P. Li, K.S. Klos, J. Lu, K.-H. Lan, Y. Nagata, D. Fang, T. Jing, D. Yu, ErbB2 promotes Src synthesis and stability: novel mechanisms of Src activation that confer breast cancer metastasis., Cancer Res. 65 (2005) 1858–67. doi:10.1158/0008-5472.CAN-04-2353.

